# The etiological classification of the epilepsies: A brain network underpinning?

**DOI:** 10.1101/739052

**Authors:** Geertruida Slinger, Willem M. Otte, Lotte Noorlag, Floor E. Jansen, Kees P.J. Braun, Eric van Diessen

## Abstract

**Objective:** the current epilepsy classification is primarily clinical driven and lacks a mechanistic basis. A mechanistic basis of the classification, and within the classification especially the etiology layer, may help to better understand epilepsy and the associated comorbidities. It may also be helpful in guiding epilepsy treatment. With this study we aimed to investigate if there is a modelled mechanistic underpinning for the etiological epilepsy classification by assessing the association between epilepsy etiology and brain network topology.

**Methods:** to that aim we assessed the association between epilepsy etiology and brain network topology. We included children referred to our outpatient first seizure clinic with suspected epilepsy who had a standard interictal EEG recording. From these EEGs, functional networks were constructed based on eyes-closed resting state time-series. Networks were characterized using measures of segregation, integration, centrality, and network strength. Principal component analyses were used to assess whether patients with epilepsy of similar etiology cluster together based on their functional brain network topology.

**Results:** in total, 228 children with epilepsy were included. Another 402 children served as control subjects. We were not able to detect a correlation between epilepsy etiology and functional brain network topology. We also did not find a difference in brain network topology between the controls and patients with epilepsy.

**Conclusions:** our results do not support the presence of a brain network underpinning for the etiological epilepsy classification. This may support the hypothesis that brain network abnormalities in epilepsy are a result of ongoing seizure activity rather than the epilepsy etiology itself. Further in-depth analyses of network measures and longitudinal studies are needed to confirm this hypothesis.

## 1. Introduction

Classification systems may help to better understand diseases with their associated signs and symptoms (Barçin and Aktekin, 2014) as they illuminate the possible analogous pathophysiological mechanisms and the involvement of different organs, tissues, and cells. Classification systems exist for most common disease or disease groups, including epilepsy. The first scheme for the classification of epileptic seizures was published by the International League Against Epilepsy (ILAE) in 1969 (Gastaut, 1969). The epilepsy classification was primarily designed for diagnosing and treating patients, but also facilitates epilepsy research and communication among clinicians worldwide (Barçin and Aktekin, 2014; Scheffer *et al*., 2017). From its inception in 1969, the epilepsy classification has been revised several times to better reflect developments in the understanding of epilepsy as well as advances in diagnostic and treatment options (Barçin and Aktekin, 2014).

The latest version of the epilepsy classification was proposed by the ILAE in 2017 (Scheffer *et al*., 2017). Although this classification aims to incorporate recent scientific insights into epilepsy, the level of knowledge is not yet sufficiently advanced for constructing an entirely scientific based classification (Scheffer *et al*., 2017). The biggest gap of knowledge – which also is the hardest to bridge – is that the fundamental neurological basis of the epilepsies is not fully understood (Scheffer *et al*., 2016). The classification system therefore continues to rely to a large extent on proxy measures including semiology, electroencephalography (EEG) features and even expert opinions. As a consequence, the epilepsy classification has a high pragmatic utility in clinical settings, but lacks a more mechanistic basis. A mechanistic underpinning of the epilepsy classification in general may be useful to better understand disease mechanisms (Manford, 2017), associated cognitive and behavioral impairments (Seidenberg, Pulsipher and Hermann, 2009; Srinivas and Shah, 2017), and the clinical course of an epilepsy, while it also supports clinical management and the development of new treatment strategies. Since the epilepsy etiology is regarded as a major determinant of the clinical course (and prognosis) and epilepsy treatment (Shorvon, 2011), we focus on the etiological epilepsy classification in this study.

A theoretical approach that may be able to capture a mechanistic underpinning of the current – primarily clinically driven – epilepsy classification is perceiving the brain as a complex network (Bullmore and Sporns, 2009). A brain network is a simplified mathematical representation of the different areas the brain consists of and how (strong) they are connected to each other. Network analysis can be applied to both structural (structural MRI) and functional (functional MRI, magnetoencephalography, EEG) brain mapping techniques, resulting in structural and functional networks, respectively (Rubinov and Sporns, 2010). Structural and functional brain networks can be characterized by a variety of local and global network measures characterizing the topological properties of the network (Rubinov and Sporns, 2010).

Previous research has extensively shown that epilepsy can be perceived as a network disorder (Kramer and Cash, 2012; van Diessen *et al*., 2013). In this study we thus aim to investigate if there is a brain network underpinning for the etiological epilepsy classification by assessing the relationship between a patient’s epilepsy etiology and the brain network configuration. We thereby focus on functional EEG based brain networks. We hypothesized that the etiological subdivision of epilepsies can also be found in the functional brain network organization, meaning there is a neural network underpinning for the current etiological epilepsy classification.

## 2. Methods

### 2.1 Patient selection

We retrospectively collected data of children (0-18 years) referred to the outpatient First Seizure Clinic (FSC) of the University Medical Center Utrecht between January 2008 and May 2018 after one or more paroxysmal event(s) suspicious for epilepsy, who had a standard EEG recording of sufficient quality. Epilepsy was diagnosed by a child neurologist based on clinical presentation, EEG findings, additional testing (for instance sleep-deprivation EEG, neuroimaging, lumbar puncture, genetic testing), and at least one year of clinical follow-up. Children in whom the diagnosis of epilepsy was rejected, were included as control group. For some children, it was still unclear whether they had epilepsy or not after additional testing and clinical follow-up. These children were excluded from our study. We also excluded children who already had an established diagnosis of epilepsy at the time of visiting the FSC and children who were on anti-epileptic drug treatment at the time of the EEG recording. We did not exclude children who were diagnosed with epilepsy in the past and presenting at the FSC after their past epilepsy was believed to be resolved. For those who visited the FSC more than once, we only included the EEG data of the first visit.

The institutional Medical Research Ethics Committee approved the use of the retrospectively collected patient data and concluded that the Medical Research Involving Human Subjects Act did not apply. The need for informed consent was waived, provided that the data were handled anonymously.

### 2.2 Data items

From all children eligible for inclusion in our study, we systematically collected the FSC EEG recording and corresponding report, final diagnosis (epilepsy or no epilepsy), and the following additional clinical characteristics: sex, age at first seizure-like event, age at EEG, developmental status, general medical history, and family history with respect to seizures. Personal data items that could directly be traced back to the included children were anonymized.

For the children diagnosed with epilepsy, we scored the epilepsy type (focal, generalized or both), the epilepsy syndrome if applicable, and the presumed epilepsy etiology (structural, genetic, metabolic, immunological, infectious or unknown) according to the most recent epilepsy classification (Scheffer *et al*., 2017). All epilepsy syndromes we considered in this study, are listed in Supplementary Table 1. Some patients’ epilepsies were classifiable in more than one etiological group. Since the etiological classification is not hierarchical, one cannot say that one etiology is more important than another. We therefore chose to score the etiology most important for the actual seizures to occur for epilepsies with more than one etiology. For instance: an epilepsy due to a focal cortical dysplasia based on a gene abnormality has both a genetic and a structural etiology, but the brain lesion is primarily responsible for seizure generation – and thus a structural etiology is scored.

We used the web-based software tool for clinical study data OpenClinica to systematically store all data. Data collection was performed by two authors [EvD and GS] separately. Difficulties in scoring the data-items were resolved by discussing them with another author [LN] and, if necessary, with a child neurologist [FJ]. Part of our dataset (children visiting the FSC between January 2008 and May 2013) was already published in a previous study (van Diessen *et al*., 2018; training cohort).

### 2.3 Data acquisition and selection

All EEGs were recorded with at least 21 scalp electrodes (Fp1, Fp2, F3, F4, F7, F8, Fz, C3, C4, Cz, T7, T8, P3, P4, P7, P8, Pz, O1, O2, A1, A2), arranged according to the international 10-20 system (SystemPLUS EVOLUTION, Micromed, Italy). Some EEG recordings also included the two additional electrodes F9 and F10. Impedance of each electrode was kept below 5 kΩ. The sampling frequency of the EEGs ranged between 512 and 2048 Hz.

For network analysis, we only needed the resting state parts of the EEGs. Quality of all EEGs was reviewed by visual inspection by two of the authors [EvD and GS]. Difficulties were resolved by discussion. Bad quality EEGs were excluded from further analyses. An EEG was assessed as being bad when at least one of the following exclusion criteria was met: EEG registration with electrode cap, missing electrodes, EEG full of artefacts (myogenic and/or electrode-contact artefacts in electrodes other than Fp1/2, A1/2, F9/10, see below), sleep or altered state of consciousness during EEG, and no or too short (< fifteen seconds) eyes-closed resting state. See also Supplementary Figure 1 for an overview of our EEG quality assessment. For all EEGs of sufficient quality, we selected an epoch of fifteen seconds interictal, eyes-closed resting state EEG, preferably without spikes and minor artefacts.

EEGs were anonymized and exported as raw Micromed files (TRC format), with standard G2 referencing. The frontoparietal (Fp1 and Fp2) and basal temporal (A1 and A2) electrodes were excluded from data analysis, since they often contain eye-movement artefacts. If present, additional channels F9 and F10 were also excluded to make sure that every brain network would contain an equal number of nodes across the group (that is: electrodes, n=17). Before plotting networks, all resting state epochs were down sampled to 256 Hz and re-referenced against the average reference electrode. Additionally, all EEG data were band-pass filtered into the common broadband (0.5-30 Hz), delta (0.5-4.0 Hz), theta (4-8 Hz), alpha (8-13 Hz), and beta (13-30 Hz) band. We excluded the gamma-band (30-80 Hz) from further analyses, as this band is prone to be contaminated with muscle artefacts. Epoch processing was carried out using MATLAB and R software.

### 2.4 Functional connectivity

Brain functional networks were constructed per fifteen second epoch, based on the functional connectivity between each pair of electrodes, resulting in a square symmetric 17 by 17 functional connectivity matrix. Functional connectivity was assessed using the phase lag index (PLI). The PLI is a value between 0 and 1 indicating the level of phase synchronization between two EEG signals. (Stam, Nolte and Daffertshofer, 2007) The higher the PLI value, the higher the level of synchronization and the stronger the functional connectivity.

### 2.5 Network construction and graph theoretical analysis

Based on the connectivity matrix, a weighted directed network was constructed for each subject, expressed with the graph *G = (V, E, W)*, consisting of *V* vertices (nodes, 17 in our study) and *E* edges (connections) and *W* representing the symmetric *V × V* connectivity matrix with *W*_*ij*_ quantifying the connection strength as determined with the PLI between vertices *v*_*i*_ and *v*_*j*_.

Functional brain network properties were described and quantified using weighted measures of segregation (clustering coefficient (C), modularity (Q)), integration (characteristic path length (L)), and centrality (betweenness centrality (BC) and closeness centrality (CC)) (Figure 1). In addition, we also calculated the average network strength. The network’s integration reflects the ability to rapidly combine information from different brain areas. Measures of segregation indicate the network’s ability of specialized processing in local circuits. Centrality measures assess the importance of the individual nodes in a network (Rubinov and Sporns, 2010). Brief descriptions of the specific metrics are provided in Supplementary Table 2. The metric formulas are published in various previous studies (Rubinov and Sporns, 2010). All metrics were calculated from the weighted adjacency matrix using the NetworkToolbox package in R.

**Figure 1.**
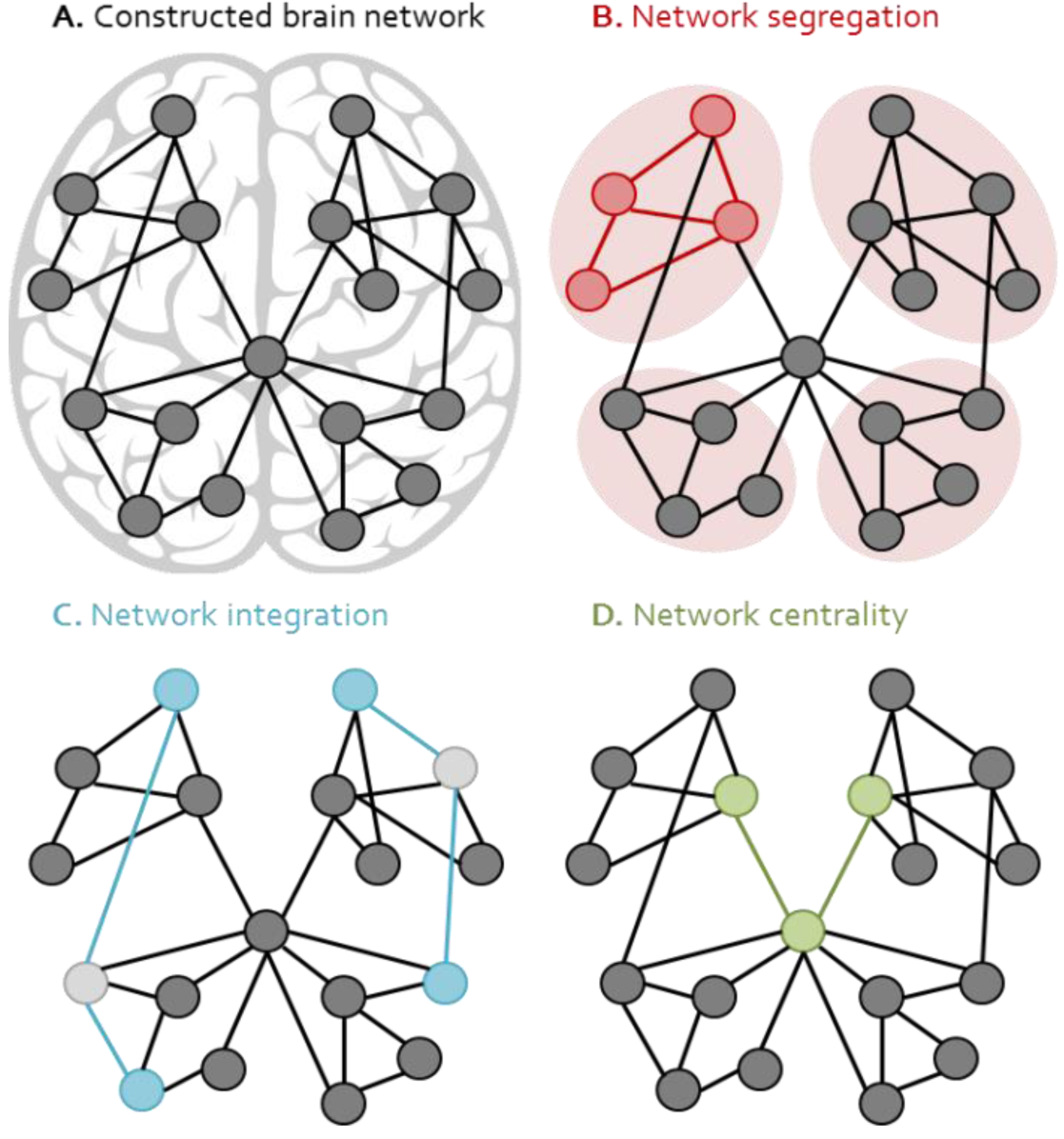
Schematic representation of network metrics. A) A constructed brain network consisting of nodes (circles) and edges (lines). B) Network segregation; the clustering coefficient is based on triangles in a network, while the modularity characterizes the modular (ellipses) structure of a network. C) Network integration; the shortest path length (in blue) is indicative for the efficiency of the exchange of information between brain areas. D) Network centrality; centrality measures identify the most central nodes in and their importance for a network.

### 2.6 Principal component analysis

We used principal component analysis (PCA) to analyze our data. PCA is a statistical tool to reduce the number of variables (that is: the dimensionality) of a dataset, while retaining as much as possible of the variation of this set (Jolliffe and Cadima, 2016). This analysis was required to deal with the relatively large number of network characteristics in our dataset. For variable reduction in a PCA, a number of original, correlated variables are transformed into a smaller number of uncorrelated variables: the principal components. So, instead of analyzing a variety of original variables, just a limited number of components containing the majority of the dataset’s variation are investigated (Groth *et al*., 2013). The first principal component always explains the highest percentage of the variability in the data, and each succeeding component explains the highest percentage of the remaining variability.

With the intention to reveal clusters of participants with it, we chose to do a PCA for a dataset split on epilepsy diagnosis (epilepsy versus no epilepsy) and for a dataset split on epilepsy etiology (only epilepsy patients). Principal component analyses were performed per frequency band. Next to the network measures, we also included gender, developmental status, and age at presentation as variables in our analyses, since it is known that these factors can influence functional network topologies, and thus can cause variability in the data (van den Heuvel *et al*., 2009; Boersma *et al*., 2011; Smit *et al*., 2012). Analyses were thus performed with a total of eleven variables for each frequency band: nine quantitative variables (maximum and medium betweenness centrality, maximum and medium closeness centrality, clustering coefficient, age at EEG recording), and two qualitative (gender and developmental status) variables. For PCAs including both quantitative (that is: numerical) and qualitative variables (that is: categorical), we used the PCAmix package in R. Since the results of a PCA depend on the scales of measurement of the included variables, we standardized the quantitative variables prior to conducting the PCAs by subtracting the variable mean of and dividing each variable data point by the variable standard deviation. Figure 2 represents a schematic overview of our methodological work-up.

**Figure 2.**
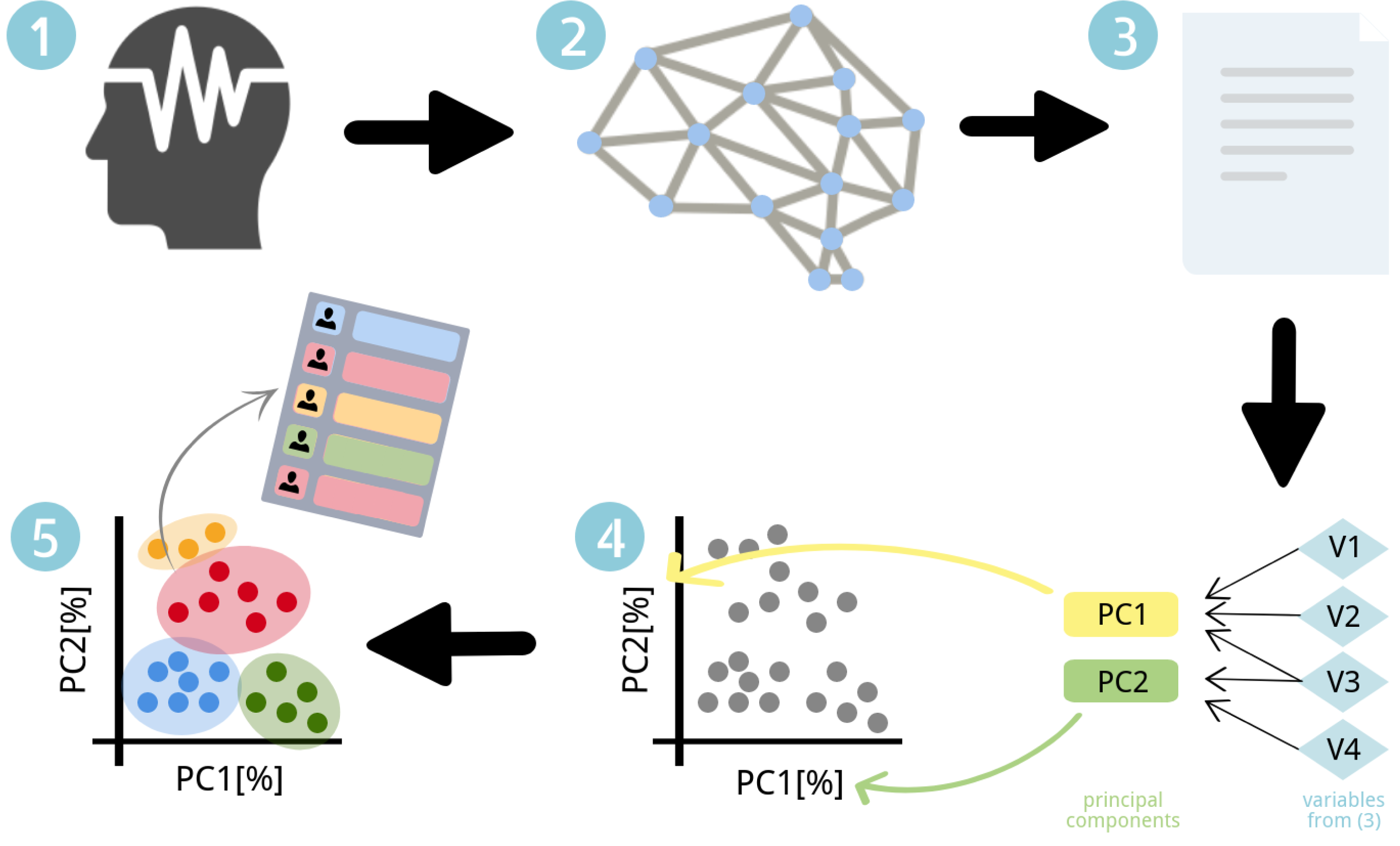
Schematic overview of methodological work-up. 1) Standard EEG recording for all participants, retrospectively collected. 2) Construction of EEG based functional brain networks. 3) An overview of calculated network measures and demographical data items form the input for the principal component analyses. 4) Principal component analyses reduce the number of correlated variables to a smaller number of uncorrelated principal components. 5) Cluster analysis detects clusters in the PCA plots, which can be related to clinical data (diagnosis, epilepsy etiology). Since there were obviously no clusters present in our data, we did not perform the cluster analysis. PC: principal component, V: variable.

## 3. Results

### 3.1 Demographics

In total, we identified 981 children who visited the FSC of the University Medical Center Utrecht between 1 January 2008 and 31 May 2018, of whom 630 were eligible for inclusion in our study (Figure 3). From these 630 children, 228 (36.2%) were diagnosed with epilepsy and the remaining 402 (63.8%) subjects were included as controls. There were no major differences in baseline characteristics between the patient and control group, except for the developmental status. A delayed development (intelligence quotient (IQ) < 70) was slightly more often seen in the patient than in the control group (21.9% versus 16.2%). All baseline characteristics for both the patients with epilepsy and the control subjects are listed in Table 1.

**Table 1.** Baseline characteristics. Baseline characteristics of the children included in our study.

|  | Epilepsy | No epilepsy | All |
| --- | --- | --- | --- |
| **Demographics** |  |  |  |
| Subjects | 228 (36.2) | 402 (63.8) | 630 (100) |
| Male gender | 131 (57.5) | 219 (54.5) | 350 (55.6) |
| Age at EEG recording | 8.6 ± 4.0 | 8.3 ± 4.5 | 8.4 ± 4.3 |
| Delayed development | 50 (21.9) | 65 (16.2) | 115 (18.2) |
| **Epilepsy etiology** |  |  |  |
| Unknown | 95 (41.7) | - | - |
| Genetic | 83 (36.4) | - | - |
| Structural | 45 (19.7) | - | - |
| Metabolic | 5 (2.2) | - | - |
| Infectious | 0 (0.0) | - | - |
| Immune | 0 (0.0) | - | - |

All numbers are displayed as *absolute number (percentage)*, except for the age at EEG recording, which is displayed as *mean ± SD*. The age is represented in *years.months*.

**Figure 3.**
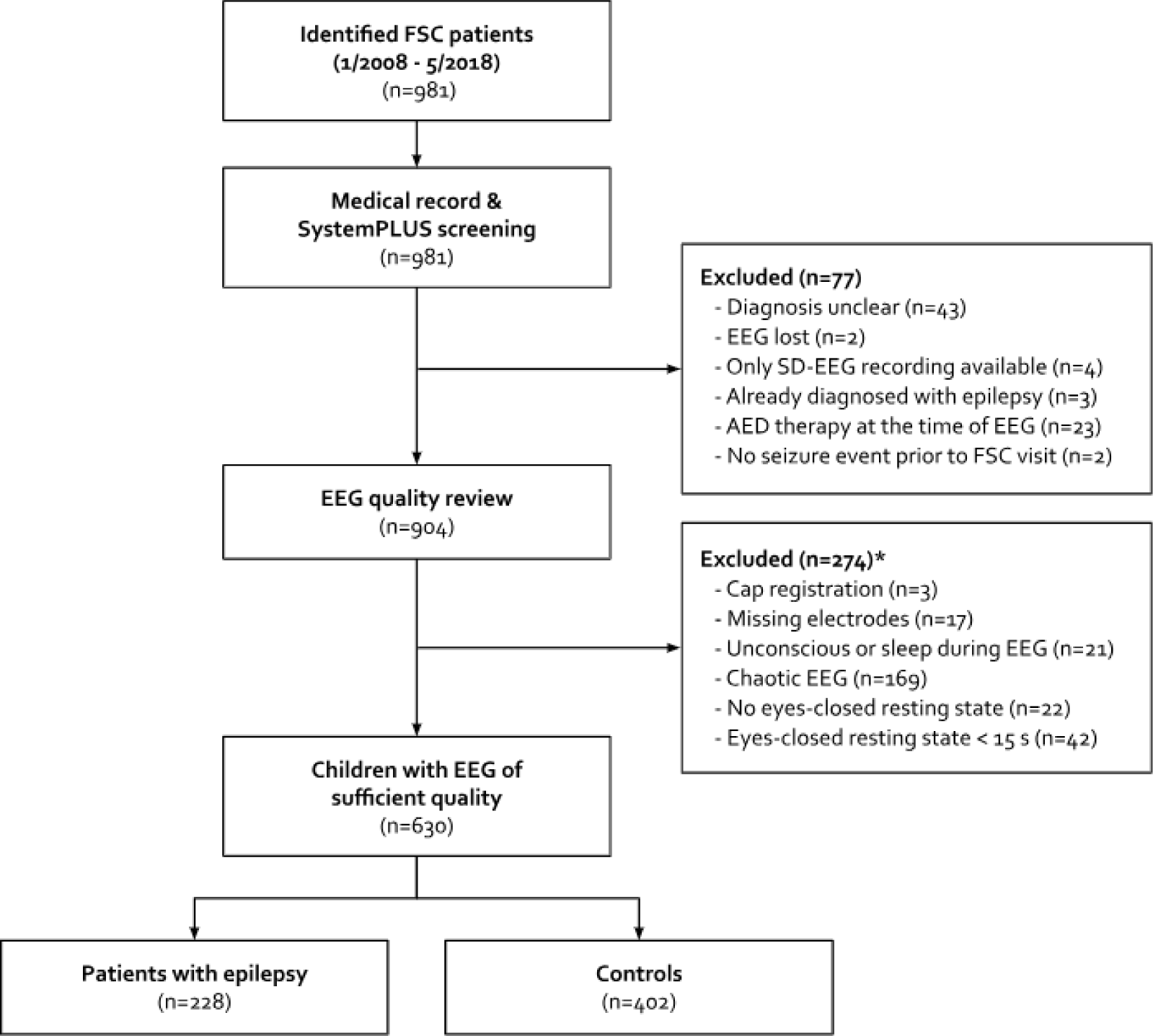
Subject flowchart. From the 981 identified children 630 were eligible for inclusion in our study. Information on final diagnosis was already collected, but not accessed during the EEG quality review process to avoid selective inclusion. *EEGs were reviewed using the review flowchart as depicted in Supplementary Figure 1. AED: anti-epileptic drug, FSC: First Seizure Clinic.

Regarding the patients’ epilepsies, an unknown etiology was most common (95 children, 41.7%). 83 children (36.4%) had an epilepsy of genetic origin and 45 children (19.7%) were diagnosed with epilepsy due to structural brain abnormalities (acquired or congenital). A metabolic etiology was considered for five (2.2%) patients’ epilepsies. Epilepsies of infectious and immune origins were not observed in our patient cohort (Table 1).

### 3.2 Principal component analyses

#### 3.2.1 Principal component analyses epilepsy versus no epilepsy

The proportion of variance explained by the first dimension or principal component ranged between 47.6% (delta band) and 54.7% (alpha band). The proportion of variance explained by the second principal component ranged between 10.4% (alpha band) and 13.1% (delta band). See Figure 4A for the broadband data. Plots for the other frequency bands can be found in Supplementary Figure 2. Variables strongly correlating (r^2^ > 0.7) with principal component 1 (and with each other), were the maximum and medium closeness centrality, the path length, and network strength. Principal component 2 only showed moderate correlations (r^2^ 0.3-0.7) with the maximum and median betweenness centrality (Figure 5A and Table 2 (broadband), Supplementary Table 3 (other bands)). With visual inspection, no clusters could be observed in any of the data plots. Therefore, no further cluster analyses have been performed.

**Table 2.** Correlations between variables and principal component 1 and 2 for the broadband.

|  | diagnosis |  | etiology |  |
| --- | --- | --- | --- | --- |
|  | PC1 | PC2 | PC1 | PC2 |
| *Numerical variables* |  |  |  |  |
| BC_max | 0.191 | 0.568 | 0.259 | 0.335 |
| BC_median | 0.391 | 0.360 | 0.456 | 0.189 |
| C | 0.917 | 0.060 | 0.920 | 0.043 |
| CC_max | 0.959 | 0.001 | 0.956 | 0.001 |
| CC_median | 0.933 | 0.041 | 0.932 | 0.033 |
| EEG_age | 0.036 | 0.081 | 0.018 | 0.228 |
| PL | 0.845 | 0.024 | 0.837 | 0.018 |
| Q | 0.445 | 0.008 | 0.479 | 0.000 |
| Strength | 0.928 | 0.055 | 0.929 | 0.041 |
| *Categorical variables* |  |  |  |  |
| Development | 0.000 | 0.010 | 0.009 | 0.315 |
| Gender | 0.004 | 0.007 | 0.000 | 0.002 |

**Figure 4.**
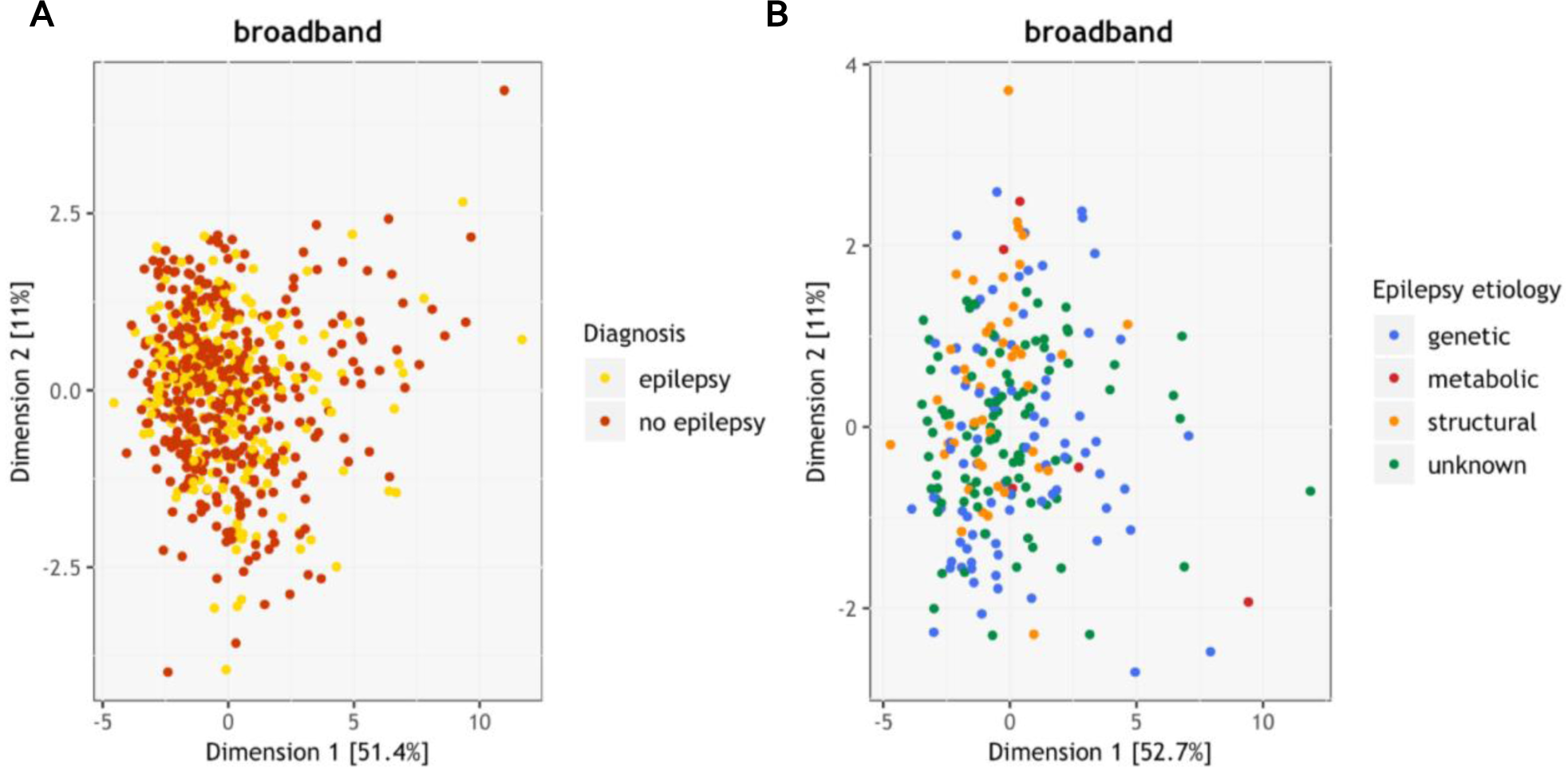
Results of principal component analyses for the broadband. A) Distribution of individuals (n=630) at PCA dimension 1 (horizontal) and 2 (vertical), split on final diagnosis. B) Distribution of individuals (n=228) with epilepsy at PCA dimension 1 (horizontal) and 2 (vertical), split on epilepsy etiology. Percentages in the brackets display the percentages of variance in the data explained by the corresponding PCA dimension.

#### 3.2.2 Principal component analyses epilepsy etiologies

The proportion of variance explained by the first dimension or principal component ranged between 46.7% (beta band) and 54.0% (alpha band). The proportion of variance explained by the second principal component ranged between 11.0% (broadband) and 12.6% (beta band). See Figure 4B for the broadband data. Plots for the other frequency bands can be found in Supplementary Figure 3. Variables strongly correlating (r^2^ > 0.7) with principal component 1 (and with each other), were the maximum and medium closeness centrality, the path length, and network strength. Principal component 2 only showed moderate correlations (r^2^ 0.3-0.7) with the maximum betweenness centrality and gender (Figure 5B and Table 2 (broadband), Supplementary Table 3 (other bands)). With visual inspection, no clusters could be observed in any of the data plots. Therefore, no further cluster analyses have been performed.

**Figure 5.**
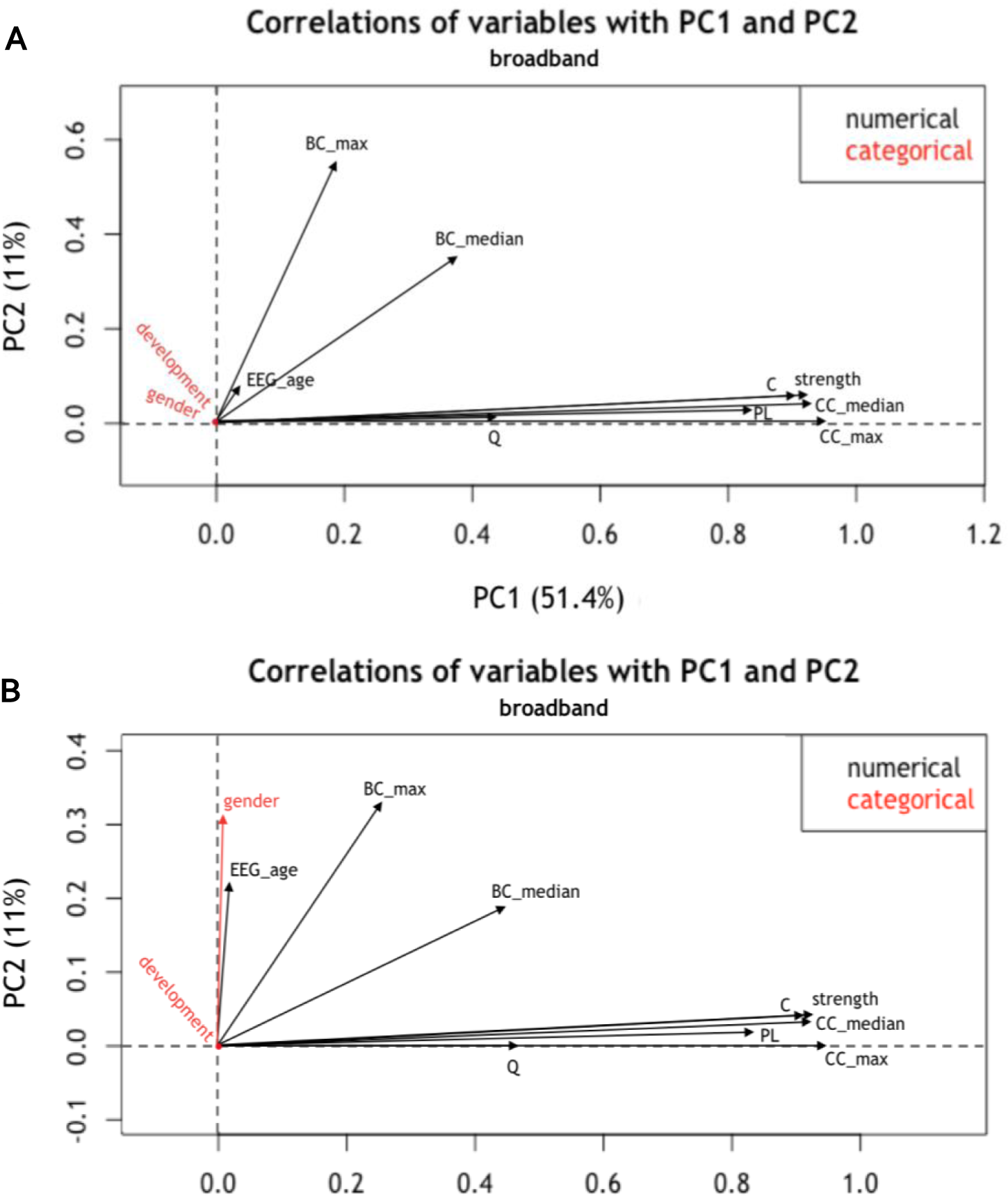
Correlation between the included variables and principal component 1 and 2 for the broadband. A) Correlation of variables with principal component 1 and 2 for the PCA epilepsy versus no epilepsy. B) Correlation of variables with principal component 1 and 2 for the PCA epilepsy etiology. BC_max: maximum betweenness centrality, BC_median: median betweenness centrality, C: clustering coefficient, CC_max: maximum closeness centrality, CC_median: median closeness centrality, EEG_age: age at EEG recording, PC: principal component, PL: characteristic path length, Q: modularity.

## 4. Discussion

In this large pediatric network study, we assessed the possible association between epilepsy etiology and global functional brain network organization. By means of principal component analyses, we were not able to detect such an association, meaning that we did not find the distinct epilepsy etiologies to be represented by specific functional brain network topologies. Besides, we were also not able to differentiate between patients with epilepsy and control subjects in the principal component analyses. These results apply for all frequency bands explored in our study (broadband, delta, theta, alpha, and beta band). Further, our results show that the average clustering coefficient, characteristic path length, closeness centrality (median and maximum), and network strength are highly correlated with each other in all frequency bands, either positively or negatively.

### Mechanistic concepts for network alteration in epilepsy

Brain functioning in its complexity is currently incompletely understood and not fully mechanistically interpretable. Nonetheless, explaining the essential brain manifestations in terms of mechanisms may help unraveling pathophysiological behavior and dysfunction. This requires developing models with some level of abstraction and simplification. Two basic models regarding network alterations in epilepsy have evolved over time. The first model states that epileptic activity itself causes changes in the brain network organization. This theory is supported by the finding that network abnormalities become more prominent as the duration of epilepsy increases (Qiu *et al*., 2017; Park *et al*., 2018). The second model focuses on the epilepsy origin primarily and states that the underlying disease etiology is directly responsible for the disruption of neural networks, which in the end may lead to seizure generation. This hypothesis does not deny the network deforming capacity of seizures but regards the epilepsy etiology as the most important determinant of network abnormalities (Scott, 2016). If the latter hypothesis holds true, one could expect that there are differences in the brain network topologies in epilepsies of different etiologies. Since we did not find such topological differences, our study does not directly support the etiology model. We were also not able to detect topological brain network differences between healthy children and children with epilepsy. This is in contrast with findings of subtle network changes in a selected subset of our study population that we previously described (van Diessen *et al*., 2016). It should, however, be mentioned that we did not look into network differences at the level of individual measures. Besides, the chosen network measures were limited to the most commonly used measures. It thus remains unclear if network differences are truly absent. In this light, further exploration of network topological differences, with more complex and less correlated measures, is needed (van Diessen *et al*., 2014). Furthermore, one could argue that the sparsity of a standard EEG recording might be insufficient to find network differences between patient groups. For example, a previous study has (indirectly) addressed the association between brain network topology and epilepsy etiology by means for functional MRI (Doucet *et al*., 2014). In temporal lobe epilepsy, a very subtle difference in functional brain network topology was found between patients with structural and non-structural epilepsy. Subanalysis in our study did not reveal differences for the structural epilepsies compared with epilepsies of other origins. Another explanation might be the construction of functional network in signal rather than source space. By denominating EEG electrodes as functional nodes, it remains questionable if information on the underlying networks can be inferred (van Diessen *et al*., 2015).

### Strengths and limitations of the study

Our study is unique with respect to the number of subjects included. It should, at the same time, be noted that our patient group was quite heterogeneous regarding epilepsy types, but at least the data collection and processing is equal for all patients, thereby reducing data variability. Furthermore, we only included drug-naïve children with epilepsy, thereby ruling out the effect of anti-epileptic drugs as potential confounder (Haneef, Levin and Chiang, 2015; van Veenendaal *et al*., 2017). Third, although we formulated a hypothesis beforehand, by choosing principal component analyses we let our data speak for themselves, while we did not select the network metrics to be used to plot clusters manually.

Although our study has some major strengths, several limitations should also be mentioned. Firstly, we aimed to evaluate brain network configurations for all six etiological classes postulated in the newest epilepsy classification (Scheffer *et al*., 2017). However, epilepsies of a structural, genetic, and unknown etiology are overrepresented in our cohort, while the epilepsies with an infectious, immune or metabolic etiology are very scarce or even absent. Though this is explained by the incidence of the different etiologies in childhood epilepsy and also applies to other, similar cohorts (Wirrell *et al*., 2011; Sokka *et al*., 2017), it hampers the generalization of our results to the entire etiological classification. Another factor limiting the generalizability is the unequal representation of the children’s ages in our study population. The majority of very young children had an artefact-full EEG or an EEG recording without a sufficient eyes-closed resting state period, for which we excluded them from further analysis. Although we included age at the time of EEG acquisition as a variable in our principal component analyses, we cannot state with certainty that our results are also applicable for the very young patient group. Secondly, as described earlier, some epilepsies were classifiable in more than one etiological class, of which we chose the most important one with respect to the development of seizures. This approach is pragmatic, but debatable. The same applies for children who visited the FSC twice. For most of these children, the diagnosis was equal both times, but some of them did not have clear clinical and/or electrophysiological signs for an epilepsy diagnosis at their first visit, while they were diagnosed with epilepsy several years later, at their second visit. This raises the question whether these children were allocated to the right group based on their first FSC visit. Lastly, though the first two principal components of our analyses account for a good sixty to sixty-five percent of the variance within our dataset, we still lost around one third of the original information in the data. Adding more principal components to our plots could have reduced the percentage of missed information, but also would have made the plots sparser and harder to interpret. We therefore think we chose the right balance between the amount of information in the data and the data interpretability.

### Conclusion and further directions

In conclusion, this study sought to provide a brain network underpinning for the etiology layer of the current epilepsy classification. Our results do not support the presence of such an underpinning, which may point in the direction that the underlying disease etiology is not directly responsible for brain network modulation and, consequently, seizure generation. Our study, however, has some important limitations. More studies are needed to further explore the possible association between brain network topology and the (etiological) subdivision of the epilepsies. To overcome the limitations of EEG based networks, we suggest a large (f)MRI studies. Further, longitudinal studies including patients with epilepsies of different etiologies are needed to shed more light on the role of both the epilepsy etiology and recurrent seizures in the development of brain network alterations.

## Acknowledgements

We would like to thank Sophie E. van Peer and Casper Dijkman for their help with collecting the EEG and clinical data.

## References

Barçin, E. and Aktekin, B. (2014) ‘State of the Art Approach to the Classification of Epileptic Seizures and Epilepsies’, Noropsikiyatri Arsivi, 51(3), pp. 189–194.

Boersma, M. et al. (2011) ‘Network analysis of resting state EEG in the developing young brain: structure comes with maturation.’, Human brain mapping, 32(3), pp. 413–25. doi: 10.1002/hbm.21030.

Bullmore, E. and Sporns, O. (2009) ‘Complex brain networks: graph theoretical analysis of structural and functional systems.’, Nature reviews. Neuroscience, 10(3), pp. 186–98. doi: 10.1038/nrn2575.

van Diessen, E. et al. (2013) ‘Functional and structural brain networks in epilepsy: What have we learned?’, Epilepsia, 54(11), pp. 1855–1865. doi: 10.1111/epi.12350.

van Diessen, E. et al. (2014) ‘Brain Network Organization in Focal Epilepsy: A Systematic Review and Meta-Analysis’, PLoS ONE, 9(12), p. e114606. doi: 10.1371/journal.pone.0114606.

van Diessen, E. et al. (2015) ‘Opportunities and methodological challenges in EEG and MEG resting state functional brain network research.’, Clinical neurophysiology: official journal of the International Federation of Clinical Neurophysiology, 126(8), pp. 1468–81. doi: 10.1016/j.clinph.2014.11.018.

van Diessen, E. et al. (2016) ‘Electroencephalography based functional networks in newly diagnosed childhood epilepsies’, Clinical Neurophysiology, 127(6), pp. 2325–2332. doi: 10.1016/j.clinph.2016.03.015.

van Diessen, E. et al. (2018) ‘A Prediction Model to Determine Childhood Epilepsy After 1 or More Paroxysmal Events.’, Pediatrics, 142(6). doi: 10.1542/peds.2018-0931.

Doucet, G. E. et al. (2014) ‘Early and Late Age of Seizure Onset have a Differential Impact on Brain Resting-State Organization in Temporal Lobe Epilepsy’, Brain Topography, 28(1), pp. 113–126. doi: 10.1007/s10548-014-0366-6.

Gastaut, H. (1969) ‘Classification of the epilepsies. Proposal for an international classification.’, Epilepsia, 10, p. Suppl:14–21. Available at: http://www.ncbi.nlm.nih.gov/pubmed/4980563.

Groth, D. et al. (2013) ‘Principal components analysis.’, Methods in molecular biology (Clifton, N.J.), 930, pp. 527–47. doi: 10.1007/978-1-62703-059-5_22.

Haneef, Z., Levin, H. S. and Chiang, S. (2015) ‘Brain Graph Topology Changes Associated with Anti-Epileptic Drug Use.’, Brain connectivity, 5(5), pp. 284–91. doi: 10.1089/brain.2014.0304.

van den Heuvel, M. P. et al. (2009) ‘Efficiency of Functional Brain Networks and Intellectual Performance’, Journal of Neuroscience, 29(23), pp. 7619–7624. doi: 10.1523/JNEUROSCI.1443-09.2009.

Jolliffe, I. T. and Cadima, J. (2016) ‘Principal component analysis: a review and recent developments.’, Philosophical transactions. Series A, Mathematical, physical, and engineering sciences, 374(2065), p. 20150202. doi: 10.1098/rsta.2015.0202.

Kramer, M. A. and Cash, S. S. (2012) ‘Epilepsy as a Disorder of Cortical Network Organization’, The Neuroscientist, 18(4), pp. 360–372. doi: 10.1177/1073858411422754.

Manford, M. (2017) ‘Recent advances in epilepsy.’, Journal of neurology, 264(8), pp. 1811–1824. doi: 10.1007/s00415-017-8394-2.

Park, K. M. et al. (2018) ‘Progressive topological disorganization of brain network in focal epilepsy’, Acta Neurologica Scandinavica, 137(4), pp. 425–431. doi: 10.1111/ane.12899.

Qiu, W. et al. (2017) ‘Disrupted topological organization of structural brain networks in childhood absence epilepsy’, Scientific Reports. Springer US, 7(1), pp. 1–10. doi: 10.1038/s41598-017-10778-0.

Rubinov, M. and Sporns, O. (2010) ‘Complex network measures of brain connectivity: Uses and interpretations’, NeuroImage. Elsevier Inc., 52(3), pp. 1059–1069. doi: 10.1016/j.neuroimage.2009.10.003.

Scheffer, I. E. et al. (2016) ‘Classification of the epilepsies: New concepts for discussion and debate-Special report of the ILAE Classification Task Force of the Commission for Classification and Terminology.’, Epilepsia open, 1(1–2), pp. 37–44. doi: 10.1002/epi4.5.

Scheffer, I. E. et al. (2017) ‘ILAE classification of the epilepsies: Position paper of the ILAE Commission for Classification and Terminology.’, Epilepsia, 58(4), pp. 512–521. doi: 10.1111/epi.13709.

Scott, R. C. (2016) ‘Network science for the identification of novel therapeutic targets in epilepsy.’, F1000Research, 5. doi: 10.12688/f1000research.8214.1.

Seidenberg, M., Pulsipher, D. T. and Hermann, B. (2009) ‘Association of epilepsy and comorbid conditions.’, Future neurology, 4(5), pp. 663–668. doi: 10.2217/fnl.09.32.

Shorvon, S. D. (2011) ‘The etiologic classification of epilepsy’, Epilepsia, 52(6), pp. 1052–1057. doi: 10.1111/j.1528-1167.2011.03041.x.

Smit, D. J. A. et al. (2012) ‘The brain matures with stronger functional connectivity and decreased randomness of its network.’, PloS one, 7(5), p. e36896. doi: 10.1371/journal.pone.0036896.

Sokka, A. et al. (2017) ‘Etiology, syndrome diagnosis, and cognition in childhood-onset epilepsy: A population-based study.’, Epilepsia open, 2(1), pp. 76–83. doi: 10.1002/epi4.12036.

Srinivas, H. V and Shah, U. (2017) ‘Comorbidities of epilepsy.’, Neurology India, 65(Supplement), pp. S18–S24. doi: 10.4103/neuroindia.NI_922_16.

Stam, C. J., Nolte, G. and Daffertshofer, A. (2007) ‘Phase lag index: Assessment of functional connectivity from multi channel EEG and MEG with diminished bias from common sources’, Human Brain Mapping, 28(11), pp. 1178–1193. doi: 10.1002/hbm.20346.

van Veenendaal, T. M. et al. (2017) ‘Chronic antiepileptic drug use and functional network efficiency: A functional magnetic resonance imaging study.’, World journal of radiology, 9(6), pp. 287–294. doi: 10.4329/wjr.v9.i6.287.

Wirrell, E. C. et al. (2011) ‘Incidence and classification of new-onset epilepsy and epilepsy syndromes in children in Olmsted County, Minnesota from 1980 to 2004: a population-based study.’, Epilepsy research, 95(1–2), pp. 110–8. doi: 10.1016/j.eplepsyres.2011.03.009.

