## Supplementary Material for "The etiological classification of the epilepsies: A brain network underpinning?"

### Supplementary figures and tables

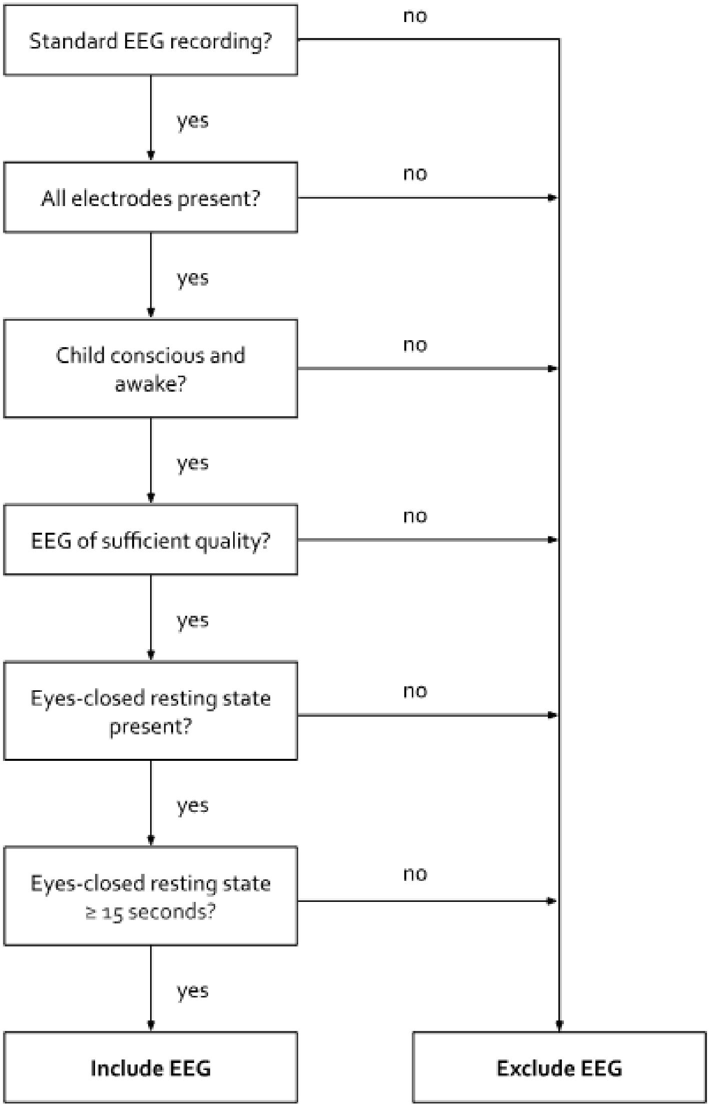


**Supplementary Figure 1 | EEG quality review flowchart.**
EEG quality was reviewed using the review flowchart. The flowchart is hierarchical, meaning that the criteria at the top are more important that the criteria at the bottom.

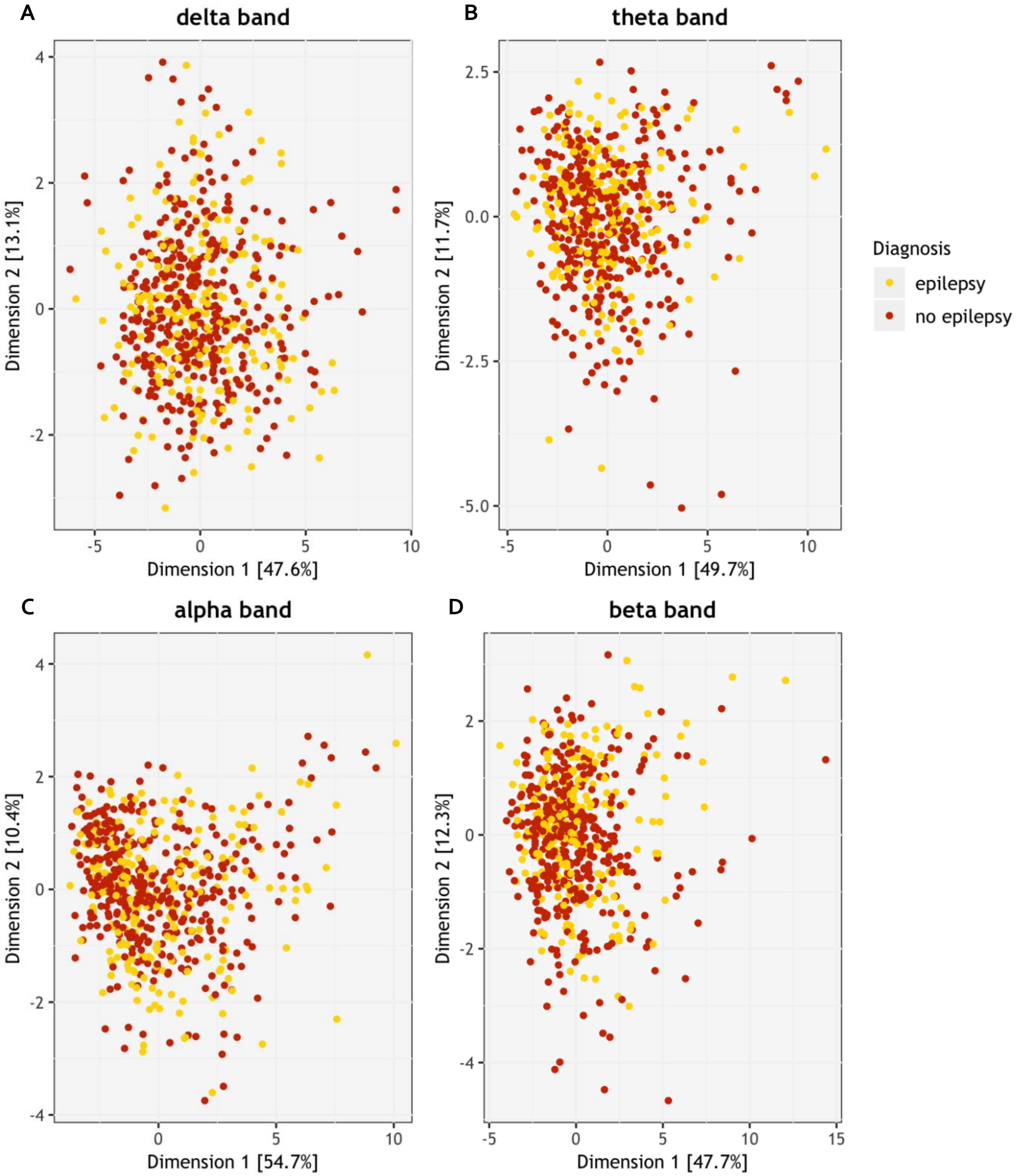


**Supplementary Figure 2 | Results of principal component analyses epilepsy versus no epilepsy**

A) Distribution of individuals at PCA dimension 1 and 2 for the delta band. B) Distribution of individuals at PCA dimension 1 and 2 for the theta band. C) Distribution of individuals at PCA dimension 1 and 2 for the alpha band. D) Distribution of individuals at PCA dimension 1 and 2 for the beta band.

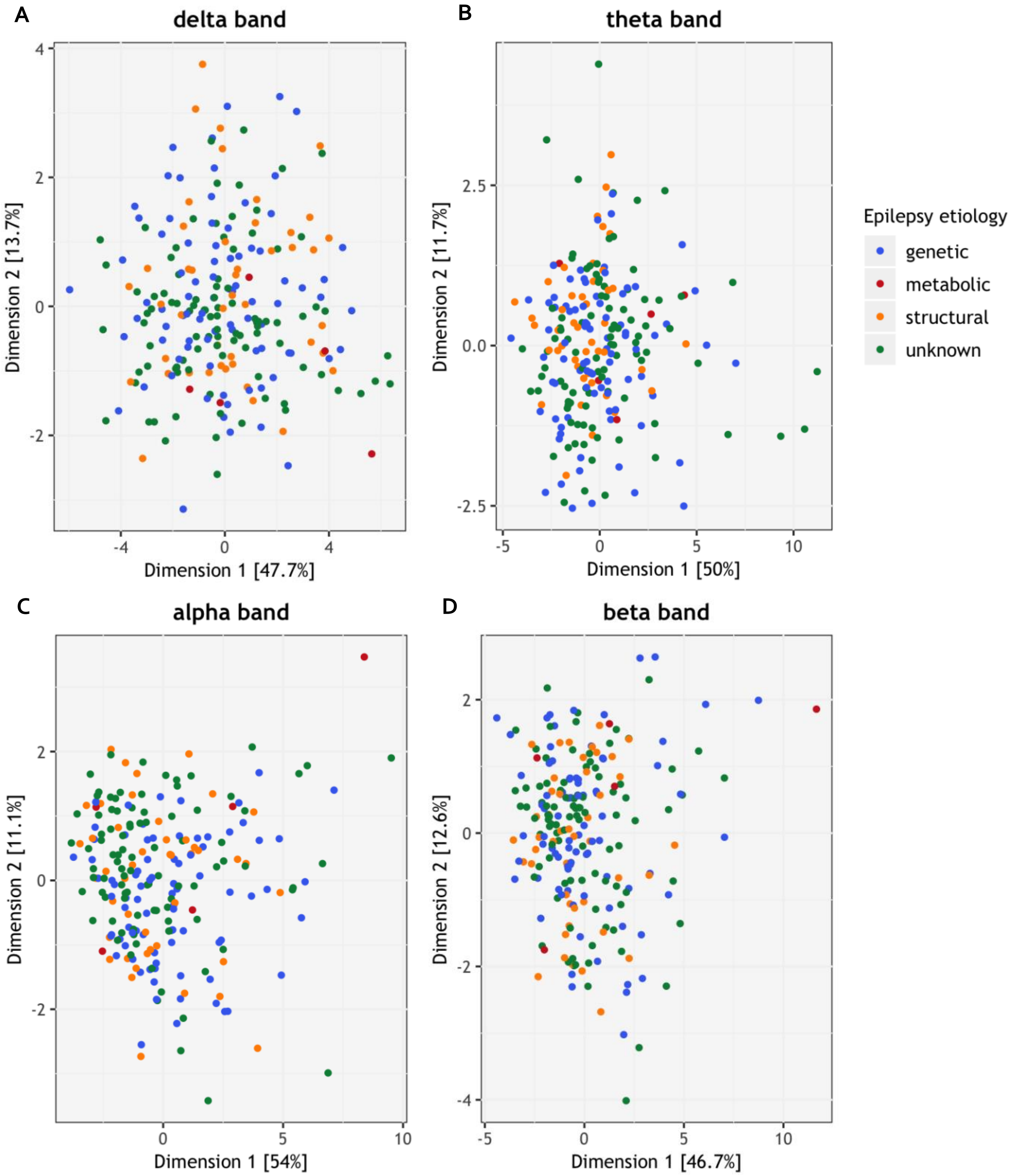


**Supplementary Figure 3 | Results of principal component analyses epilepsy etiology**
A) Distribution of individuals at PCA dimension 1 and 2 for the delta band. B) Distribution of individuals at PCA dimension 1 and 2 for the theta band. C) Distribution of individuals at PCA dimension 1 and 2 for the alpha band. D) Distribution of individuals at PCA dimension 1 and 2 for the beta band.

**Supplementary Table 1 | Epilepsy syndromes.**

| **Epilepsy syndromes by age category** |
|---|
| *Neonatal & infantile* |
| Self-limited neonatal seizures and self-limited familial neonatal epilepsy |
| Self-limited familial and non-familial infantile epilepsy |
| Early myoclonic encephalopathy |
| Ohtahara syndrome |
| West syndrome |
| Dravet syndrome |
| Myoclonic epilepsy of infancy* |
| Epilepsy of infancy with migrating focal seizures |
| Myoclonic encephalopathy in non-progressive disorders |
| Febrile seizures plus and genetic epilepsy with febrile seizures plus |
| *Childhood* |
| Epilepsy with myoclonic-atonic seizures* |
| Epilepsy with eyelid myoclonias* |
| Lennox-Gastaut syndrome |
| Childhood absence epilepsy* |
| Epilepsy with myoclonic absences* |
| Panayiotopoulos syndrome |
| Childhood occipital epilepsy (Gastaut type) |
| Photosensitive occipital lobe epilepsy |
| Childhood epilepsy with centrotemporal spikes |
| Atypical childhood epilepsy with centrotemporal spikes |
| Epileptic encephalopathy with continuous spike-and-wave during sleep |
| Landau-Kleffner syndrome |
| Autosomal dominant nocturnal frontal lobe epilepsy |
| *Adolescent & adult* |
| Juvenile absence epilepsy* |
| Juvenile myoclonic epilepsy* |
| Epilepsy with generalized tonic-clonic seizures alone* |
| Autosomal dominant epilepsy with auditory features |
| Other familial temporal lobe epilepsies |
| *Variable age* |
| Familial focal epilepsy with variable foci |
| Reflex epilepsies |
| Progressive myoclonus epilepsies |

We considered familial and idiopathic generalized epilepsy syndromes (*) to have a genetic origin. For Panayiotopoulos syndrome and (atypical) childhood epilepsy with centrotemporal spikes we scored and unknown etiology.

**Supplementary Table 2 | Definitions of the network metrics used in our study.**

| Measure | Definition |
|---|---|
| *General measures* | |
| Strength | The average sum of weight of links connected to a node. |
| *Measures of integration* | |
| Characteristic path length | The average shortest path length for all node-node connections in a network. |
| *Measures of segregation* | |
| Clustering coefficient | The fraction of a node's neighbors that are also neighbors of each other. |
| Modularity | The degree to which a network is subdivided into distinct modules, which are clusters in which the nodes are more densely connected to each other than to the other nodes in a network. |
| *Measures of centrality* | |
| Betweenness centrality | The extent to which a node lies on the shortest path between all node-node connections in a network. |
| Closeness centrality | The inverse of the average shortest path length between a node and all other nodes in a network. |

**Supplementary Table 3 | Correlations between variables and principal component 1 and 2 for PCA epilepsy versus no epilepsy (upper) panel and epilepsy etiology (lower panel)**

| | PC1 | PC2 | PC1 | PC2 | PC1 | PC2 | PC1 | PC2 |
|---|---|---|---|---|---|---|---|---|
| *Numerical variables* | *delta band* | | *theta band* | | *alpha band* | | *beta band* | |
| BC_max | 0.027 | 0.730 | 0.106 | 0.686 | 0.153 | 0.605 | 0.130 | 0.618 |
| BC_median | 0.190 | 0.496 | 0.311 | 0.414 | 0.452 | 0.230 | 0.240 | 0.522 |
| C | 0.929 | 0.038 | 0.922 | 0.054 | 0.911 | 0.055 | 0.926 | 0.047 |
| CC_max | 0.914 | 0.024 | 0.956 | 0.003 | 0.953 | 0.001 | 0.954 | 0.002 |
| CC_median | 0.936 | 0.024 | 0.938 | 0.032 | 0.938 | 0.028 | 0.939 | 0.037 |
| EEG_age | 0.018 | 0.040 | 0.054 | 0.004 | 0.227 | 0.001 | 0.087 | 0.002 |
| PL | 0.899 | 0.021 | 0.890 | 0.015 | 0.863 | 0.003 | 0.876 | 0.025 |
| Q | 0.363 | 0.009 | 0.346 | 0.011 | 0.601 | 0.003 | 0.148 | 0.052 |
| Strength | 0.946 | 0.039 | 0.933 | 0.050 | 0.921 | 0.049 | 0.941 | 0.047 |
| *Categorical variables* | | | | | | | | |
| Development | 0.003 | 0.012 | 0.004 | 0.014 | 0.000 | 0.155 | 0.000 | 0.001 |
| Gender | 0.007 | 0.012 | 0.011 | 0.008 | 0.003 | 0.019 | 0.008 | 0.000 |

| | PC1 | PC2 | PC1 | PC2 | PC1 | PC2 | PC1 | PC2 |
|---|---|---|---|---|---|---|---|---|
| *Numerical variables* | *delta band* | | *theta band* | | *alpha band* | | *beta band* | |
| BC_max | 0.005 | 0.739 | 0.115 | 0.439 | 0.093 | 0.656 | 0.067 | 0.724 |
| BC_median | 0.185 | 0.494 | 0.324 | 0.351 | 0.441 | 0.189 | 0.167 | 0.515 |
| C | 0.936 | 0.030 | 0.936 | 0.029 | 0.919 | 0.040 | 0.935 | 0.035 |
| CC_max | 0.903 | 0.029 | 0.958 | 0.000 | 0.942 | 0.001 | 0.962 | 0.007 |
| CC_median | 0.934 | 0.025 | 0.940 | 0.027 | 0.942 | 0.014 | 0.947 | 0.021 |
| EEG_age | 0.021 | 0.061 | 0.061 | 0.218 | 0.173 | 0.007 | 0.039 | 0.000 |
| PL | 0.915 | 0.012 | 0.867 | 0.017 | 0.868 | 0.000 | 0.892 | 0.003 |
| Q | 0.388 | 0.008 | 0.318 | 0.014 | 0.618 | 0.001 | 0.157 | 0.002 |
| Strength | 0.950 | 0.033 | 0.947 | 0.028 | 0.926 | 0.035 | 0.953 | 0.029 |
| *Categorical variables* | | | | | | | | |
| Development | 0.000 | 0.067 | 0.008 | 0.003 | 0.000 | 0.081 | 0.002 | 0.005 |
| Gender | 0.012 | 0.009 | 0.031 | 0.163 | 0.015 | 0.192 | 0.019 | 0.050 |

Numbers for the numerical variables are the squared correlations between the variables and the principal components. For the categorical variables, the numbers display the correlation ratios between the variables and the principal components. A correlation coefficient between 0 and 0.3 was considered weak (red), between 0.3 and 0.7 moderate (blue), and between 0.7 and 1.0 strong (green). BC_max: maximum betweenness centrality, BC_median: median betweenness centrality, C: clustering coefficient, CC_max: maximum closeness centrality, CC_median: median closeness centrality, EEG_age: age at EEG recording, PC: principal component, PL: characteristic path length, Q: modularity.
